# A low cost, open source Turbidostat design for *in-vivo* control experiments in Synthetic Biology

**DOI:** 10.1101/617423

**Authors:** Agostino Guarino, Barbara Shannon, Lucia Marucci, Claire Grierson, Nigel Savery, Mario di Bernardo

## Abstract

To characterise the dynamics of new engineered systems in Synthetic biology, continuous culture platforms are required. In this paper, after a brief review of the existing machines present in literature, we describe the design and the implementation of a new flexible and low cost turbidostat for *in-vivo* control experiments. Then, the results of a 3 hours long experiment of control of the Optical Density is reported. Since the foundation of our design is flexibility, in this work we also discuss some possible extensions of our design, with particular attention to their application to validate *in-vivo* multicellular control design.

## 1. INTRODUCTION

Biological advances over recent years have enabled the synthetic design and implementation of egineered bacterial systems to increase at an exponential rate. However, characterisation of the dynamic behaviour of these systems still proves difficult, creating a bottleneck for further advances (Mutalik et al. (2013)). Traditionally, due to ease of propagation and scalability, batch culture techniques have been used to study and characterise synthetic bacterial systems. However, it is now widely accepted that such techniques are no longer reliable. This is due to continually changing chemical environments caused by cell growth, nutrient depletion, waste production and cell death (Monod (1949)), which can lead to mischaracterisation of the output of synthetic systems in response to input.

In order to mitigate problems arising from batch culture, several continuous culture platforms have been commercially developed [Saldanha et al. (2004); Tomson et al. (2006); Lovitt and Wimpenny (1981)]. These systems add fresh media to the culture to dilute accumulated cells and waste products and are generally split into two categories, chemostats and turbidostats. Chemostats maintain bacterial cultures at steady state by performing dilution of growth media containing fresh nutrients at a fixed rate. Turbidostats, on the other hand, continually monitor the cell density of a culture via a feedback control loop and add fresh growth media when the Optical Density of the culture rises above a specified value. For characterisation of synthetic systems where cells need to be maintained at their maximum growth rate, turbidostats are generally used in preference to chemostats [Matteau et al. (2015)]. However, commercially available turbidostat systems are expensive and often lack any flexibility in design. This lack of flexibility makes them difficult to adapt to changing experimental needs; as a result many research groups have begun to develop their own designs [Hoffmann et al. (2017)].

Here we present a low-cost turbidostat design made up of modular components. This modularity allows for customisable experiment set up. Our design has the potential to be adapted and expanded for multicellular control experiments [see Fiore et al. (2017)], for which currently there are very limited options both commercially and otherwise. Our base model can be produced for less than $200, with the cost of additional components being low. Our design consists primarily of 3D printed parts which can be produced using commercially available, widely used 3D printers.

## 2. OVERVIEW OF SOME EXISTING TURBIDOSTAT IMPLEMENTATIONS

Due to the high cost of commercially available turbidostats and the increasing availability of 3D printing, several low-cost turbidostat designs have been developed by academic groups. One of the first major low-cost designs was developed by the Klavins laboratory at the University of Washington, USA [Takahashi et al. (2015)]. Klavins group designed an open-source multiplexed turbidostat system, capable of running up to eight parallel culture chambers, whose design is considered a milestone and inspired other implementations. Turbidity was measured with the use of 650 nm laser diodes and photosensors placed either side of each culture vessel. Each turbidostat housed its own electronics allowing for individual turbidity measurements to be taken. By utilising a multiplexed syringe pump system, Klavins group reduced the need for eight syringe pumps down to just a single pump, whilst still enabling individual chambers to be supplied with fresh media when required. The total cost of the build of this open source design was under $2000.

In 2016, the Khammash laboratory at ETH Zurich, proposed a turbidostat inspired by that of the Klavins lab. They added additional optogenetic features to enable the regulation of GFP production in *Escherichia coli* based on an integrated optogenetics system (Milias-Argeitis et al. (2016)). GFP production from *Escherichia coli*, dependent on the ratio of green to red light, was controlled by integral feedback. This was accomplished by adding an automated sampling system to connect the turbidostat chamber to a FACS machine. Real-time FACS analysis was able to quantify fluorescence profiles from the cells and feedback to a controller which determined the ratios of green to red light provided to the culture chamber. This design implemented 950 nm laser diode to prevent overlap in the wavelengths absorbed by the photosensitive proteins within *Escherichia coli*. In contrast to the Klavins design, this design utilised peristaltic pumps for the dilution of culture media and the automated sampling process and was somehow optimised to run optogenetic control experiments.

A different, more complex, turbidostat design was patented in Herrero et al. (2005). Therein, differently from the approaches described above, sensing of the Optical Density is obtained via the comparison of the light absorbance of two different test tubes; the first tube hosts the controlled solution, while the second a static and constant portion of growth medium that works as a constant blank reference for the evaluation of the Optical Density. While complicating the overall design, this solution allows online calibration of the OD measure but requires, higher implementation costs.

The aim of the turbidostat design presented in this paper is to have a low cost, modular and flexible machine that can be easily adapted to a different range of experiments and extended to more complex designs such as those required for the implementation of multicellular control experiments.

During the preparation of this manuscript, we became aware of another turbidostat implementation which was recently presented in McGeachy et al. (2018). Sharing a similar goal to ours, the design presented therein requires minimal 3D printed structure and electronics that can be implemented on commercial boards based on ATMEL processors.

## 3. TURBIDOSTAT DESIGN AND IMPLEMENTATION

From a control engineering viewpoint, the turbidostat schematic can be summarised as reported in figure 1a. Here, the vessel hosting the population of bacteria is considered as the system to control or “plant”; the measure of its state (Optical Density) is sensed via a light sensor and fed to the controller, which, given the reference, evaluates the control input and drives the pump (actuator). In practice, to implement this schematic, as shown in the diagram reported in figure 1b, the following items are needed:

- A media tank that stores the reservoir of culture media and supplies the pump.
- An incubator that, ideally, encases the reaction chamber and the media tank. Its presence is required in order to keep the temperature constant at 37°C and let the bacteria replicate optimally.
- A waste bottle, to collect the amount of solution pushed out of the chamber where the cells are hosted.
- In our design, it also acts as sampling outlet.
- An air pump that supplies air to the chamber to (i) push the exceeding solution out of the test tube, directly into the waste bottle; and (ii) guarantee sufficient bacteria aeration.
- A motor to continuously stir the solution via a magnet placed within the host chamber, avoiding the bacteria to aggregate and making the Optical Density homogeneous in the entire solution.

**Fig. 1.**
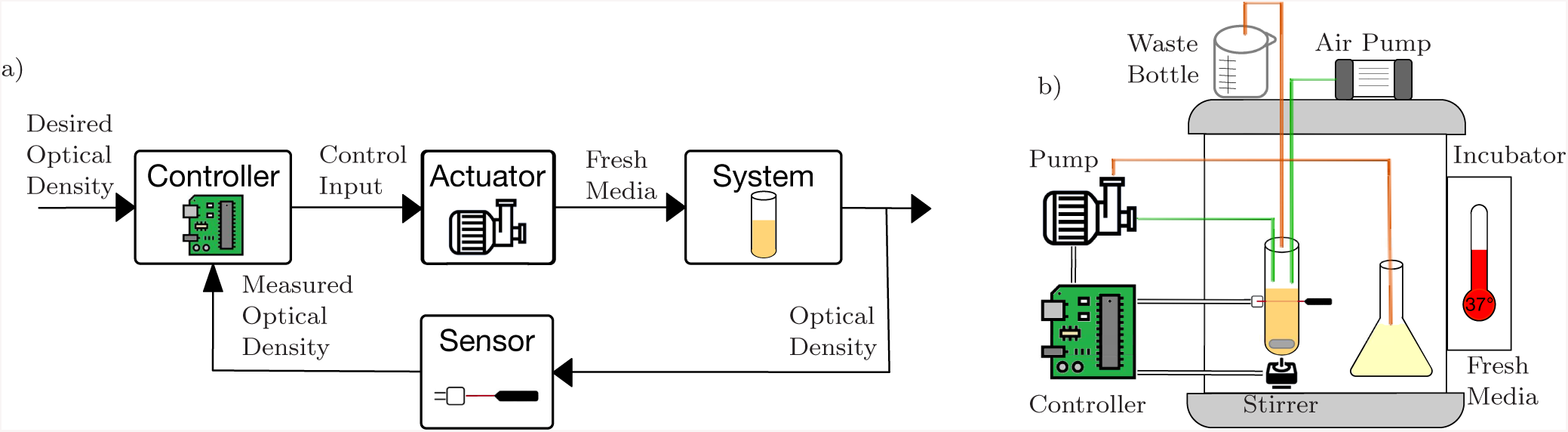
Schematic Representation of the Turbidostat. a) Control Engineering schematic of the Turbidostat control loop. b) Schematic implementation of the machine. The test tube that hosts the population of bacteria is magnetically connected to a motor that stirs the solution. The bacteria and the fresh media are placed inside an incubator to keep the temperature at 37°, to maximize bacteria’s growth rate. The air pump creates a gap of pressure between the chamber and the waste bottle: exceeding solution is pushed out when new fresh media is injected in the chamber.

In our design, all the parts to assemble the devices listed above can be 3D printed. The CAD models were redesigned and adapted so as to be parametric: in this way, beside the inner flexibility in the electronics, we guaranteed also the mechanics adaptability to a wide range of test tubes with different working volumes. The chamber support, figure 2, is inspired to Klavin’s turbidostat; however, the stirrer slot has been adapted to the most appropriate DC motor and a sliding support for the sensing circuit has been introduced. In doing this, is it possible to regulate the height at which the measure is taken, avoiding problems caused by the presence of air bubbles in the solution.

**Fig. 2.**
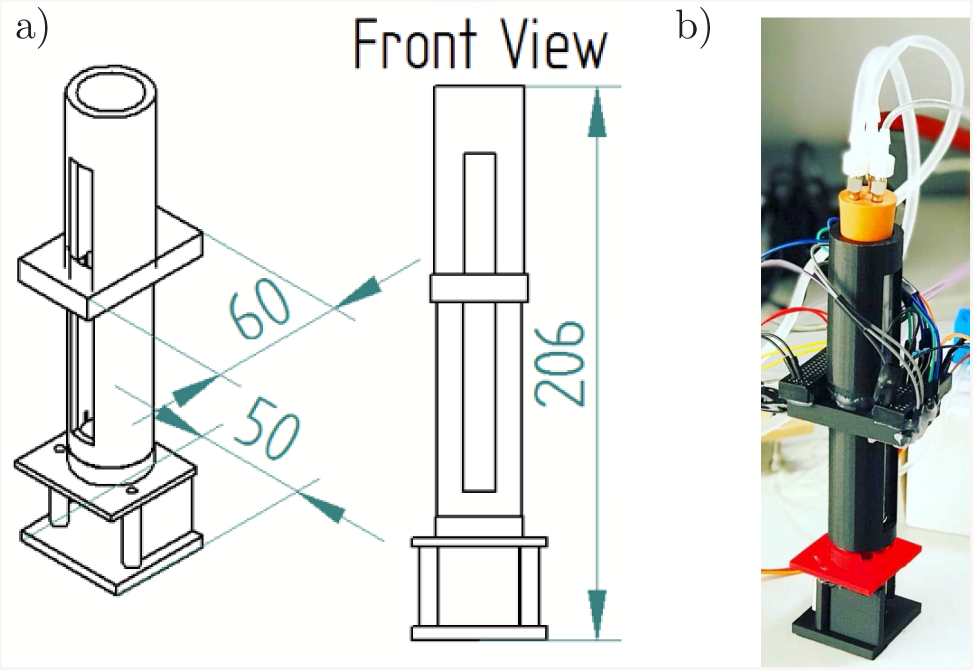
3D printed Chamber support. a) CAD model of the chamber support. Size (reported in mm) is optimised for PYREX Rimless 25×200mm test tubes. b) Assembled chamber support, with the stirrer motor mounted at the bottom, prototype of the Optical Density circuit and tube connections at the top.

For the actuators, our modular design has been tested with two different actuation systems, both of them realised via 3D printing. The first solution, 3a)-b), is to use a peristaltic pump adapted to be driven by a stepper motor. This pump is easy to drive and responds quickly but is not robust to pressure variations that might result in a backward flow. The second solution we adopted is the combination of an actuated syringe pump, shown in figure 3c), and a mutually exclusive two-ways 3D printed pinch valve, shown in figure 3d). Its driving requires the coordinated work of two servo motors, leading to longer actuation times but better robustness to pressure changes.

**Fig. 3.**
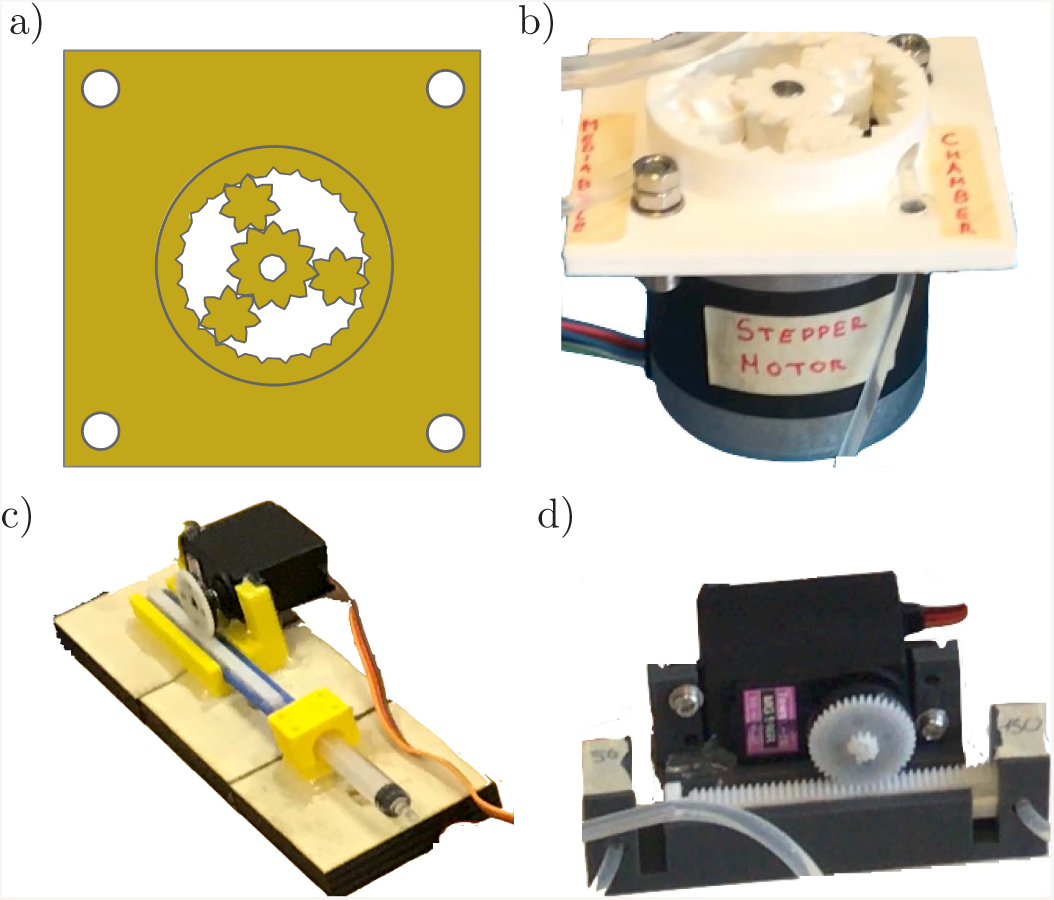
Actuation Systems. a) 3D model of the Peristaltic Pump; top view. b) Peristaltic pump driven by a stepper motor: A 4mm diameter tube passes inside the rack and it is pressed on the structure by the moving gears. During the rotation of the gears, they create a gap of pressure inside the plastic tube, moving the pressure point, and creating a flow in the same direction of rotation. c) 3D printed syringe pump. The syringe is pushed and pulled via a gearpinion mechanism driven by a servo. d) Mutually exclusive 2-ways 3D printed pinch valve. A gearpinion mechanism is moved with a servo. In the configuration shown in this panel, the way on the left is open while the one on the right is closed.

Optical Density is sensed using a photodiode in a reverse biased configuration, see figure 4. In this working condition, the diode behaves as a light-controlled current source: the current flows through a resistor of a fixed and known resistance and thus can be converted into a voltage. The higher is the Optical Density, the lower amount of light is transmitted by the solution and sensed by the photodiode; so the lower is the current and, thus, the voltage. The result is an inverse proportional relationship between the Optical Density and the voltage. Calibration experiments to reconstruct this relationship have been conducted and are reported below.

**Fig. 4.**
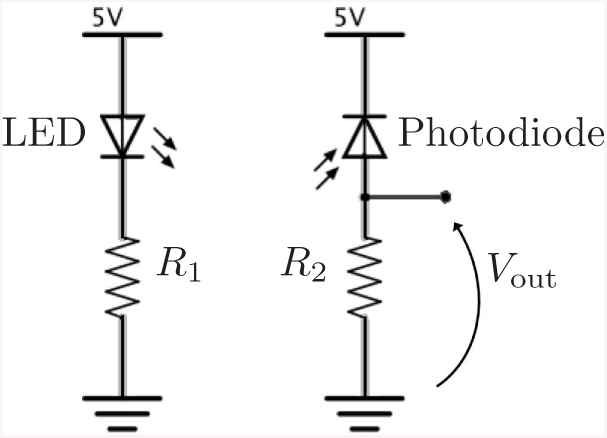
Optical Density measure circuit. On the left, a LED diode (OSRAM SHF 4544) is connected to a 5V source through the resistor *R*_1_. On the right, a photodiode (Vishay Semi-conductors BP104) is connected in a reverse bias configuration; it behaves as a light controlled current source.

The controller has been designed and implemented on an Arduino MEGA 2560 R3 board, equipped with a real-time clock module. This solution guarantees a good compromise between computing power, memory and device management compatibility. The control strategy is a classical Proportional-Integral Control, whose P and I gains were tuned heuristically depending on the type of the actuation and the working volume of the experiment.

In order to assemble a self-governing machine, a display was included in the design to provide an interactive user interface and output the current status of the machine and the instantaneous state of the experiment. The setup of the experiment can be done through a keypad through which it is possible to input several experimental parameters and the desired Optical Density. The data of the experiment is continuously logged to a common SD card that simplifies the process of data acquisition and analysis.

## 4. CALIBRATION AND VALIDATION

*Calibration experiments* were conducted to reconstruct the relationship between voltage and Optical Density. In particular, twenty-five voltage measurements of five different samples (whose OD was measured independently using a spectrophotometer - WPA CO 8000 Cell Density Meter) were performed. The test samples, together with the results of the experiment, are reported in table 1. A 2^*nd*^-order polynomial was used to interpolate the data points and obtain the voltage-OD relationship; the result is reported in figure 5. To validate the interpolated curve, the mean OD readings of the turbidostat were compared to those measured by the spectrophotometer for samples with ODs different from those used for calibration. The results are reported in table 2 confirming that the calibration is extremely precise within the range [0.05, 0.20] that is our typical working condition, while the performance worsens away from these values.

**Table 1.**
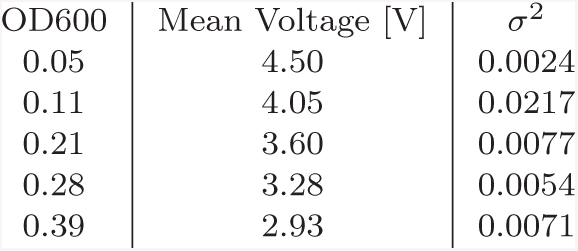
OD calibration Table.

| OD600 | Mean Voltage [V] | $\sigma^2$ |
| --- | --- | --- |
| 0.05 | 4.50 | 0.0024 |
| 0.11 | 4.05 | 0.0217 |
| 0.21 | 3.60 | 0.0077 |
| 0.28 | 3.28 | 0.0054 |
| 0.39 | 2.93 | 0.0071 |

**Table 2.** OD Validation Table.

| OD600 | Mean Read OD | Mean Reading [V] |
| --- | --- | --- |
| 0.07 | 0.0597 | 4.40 |
| 0.12 | 0.1053 | 4.09 |
| 0.15 | 0.1366 | 3.92 |
| 0.26 | 0.3174 | 3.16 |
| 0.37 | 0.4522 | 2.73 |

**Fig. 5.**
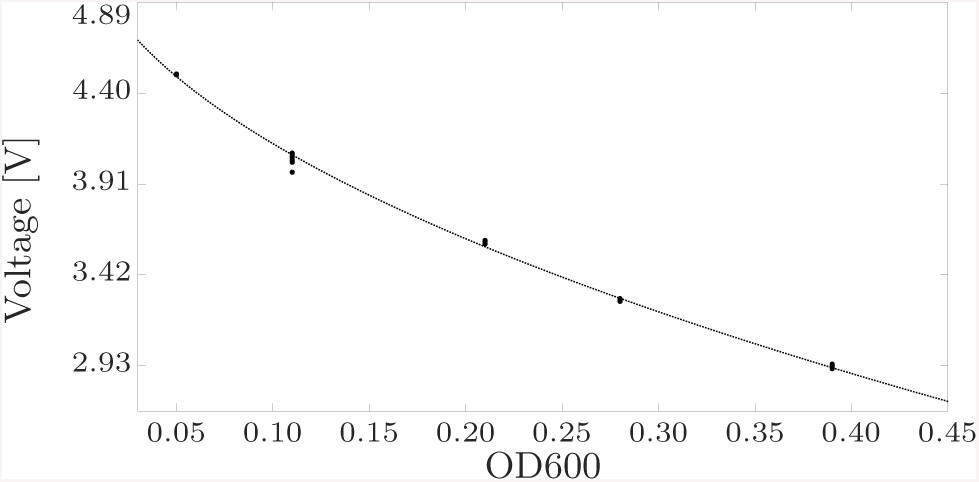
Calibration of the OD reading. Black dots represent the value of the voltage against the OD for 5 different samples. Dotted line is a 2^*nd*^ order polynomial obtained from data fitting. Polynomial coefficients are *a*_0_ ≈ 1.88, *a*_1_ ≈ −3.43 · 10^−3^, *a*_2_ ≈ 1.56 · 10^−6^.

In order to test our design, we conducted a 3 hour long closed-loop *control experiment*. As reported in figure 6, the goal was to control the OD of a solution, whose initial OD was 0.04, to the desired value of 0.10. During the first phase, which lasted about 35 minutes, the controller allowed the Optical Density to grow until the setpoint. Then the regulation phase began and an amount of fresh media, evaluated on the difference between measure and the desired value, was supplied to the solution. For the entire duration of this phase, the Optical Density falls within the acceptable range [0.09, 0.11], with an observed peak of 0.12 that might be the result of measurement noise.

**Fig. 6.**
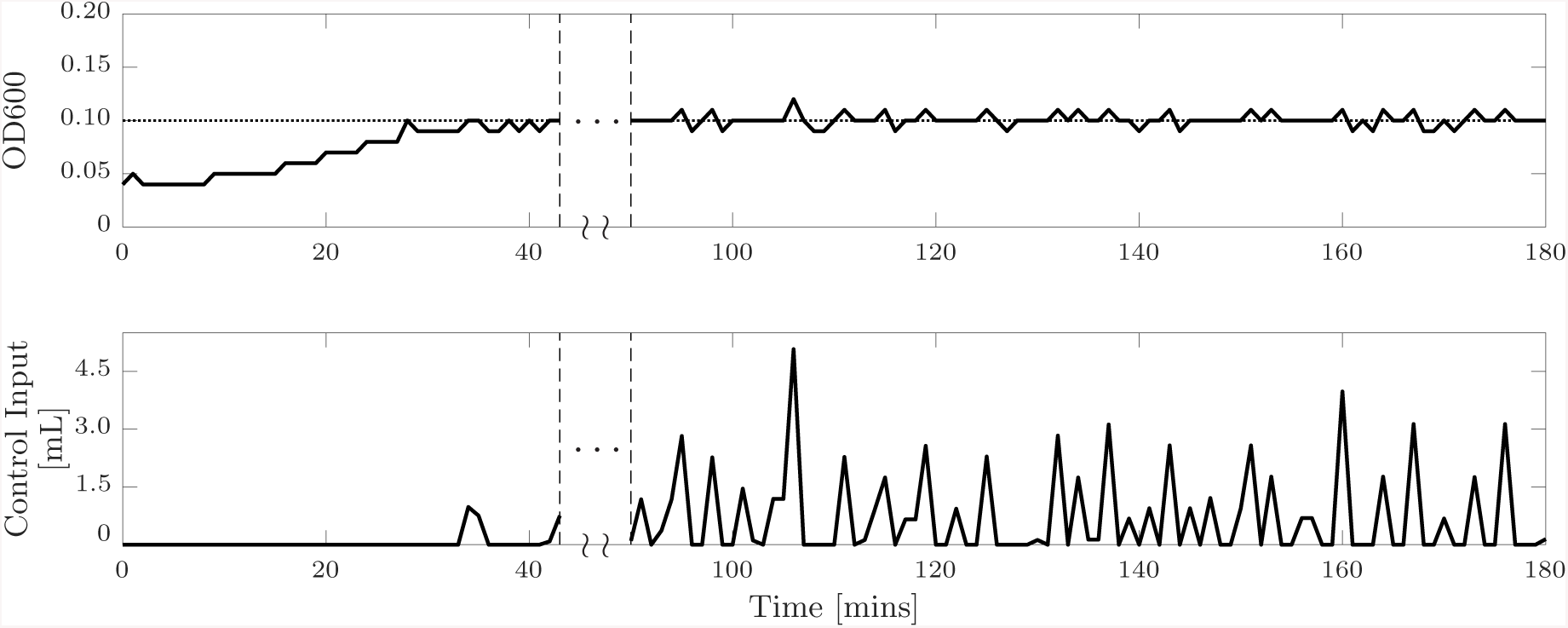
OD regulation Control experiment. Duration of the experiment : 3 hours. The setpoint of the experiment is 0.10; initial value of the OD is 0.05. During the first 30 minutes, the solution is left growing until it reaches the setpoint; then the regulation phase begins. Then, 90 minutes of regulation have been reported. The PI controller gains are *K*_p_ = 30 and *K*_i_ = 1. Sampling Time *T*_S_ = 1 min.

## 5. CONCLUSIONS

Here we have presented the design of a low cost turbidostat that was developed to maximise flexibility and ease of implementation. All the parts needed to assemble the turbidostat can be easily 3D printed and assembled (all CAD models will be provided in a dedicated website to be made available with the final version of this manuscript). Using an Arduino micro-controller, a peristaltic or syringe pump, and photodiode in a reverse bias configuration, the design we presented is self-contained and can be easily driven by a provided user interface equipped with a display and a keyboard. Closed-loop control of the OD is achieved via a PI controller tuned heuristically. Calibration and control experiments were performed and were described in the paper confirming the effectiveness of our design.

The design is simple and modular enough to be apt for adaptation and extension for use in different control setups. For example, optogenetics, which requires the use of LEDs with fixed wavelength (red 650nm, green 535nm), does not interfere with the Optical Density sensing circuit, since it works at 950nm.

Ongoing work is aimed at adapting the design to a multichamber implementation that can be used for Multicellular Control experiments. These experiments are characterised by the presence of two different populations of bacteria, one controlling a phenotype of the other. In order to maintain a desired ratio of the two populations, a key requirement for control to work, as discussed in Fiore et al. (2017),is that the two populations must be kept physically separated, while they exchange sensing molecules. A simple schematic of the design we are currently developing is reported in figure 7. The design is inspired to the structure of a microbial fuel cell [Min et al. (2005)] where two different chambers are connected to exchange molecules through a nanoporose membrane. We are currently working on such an extension that will be published elsewhere.

**Fig. 7.**
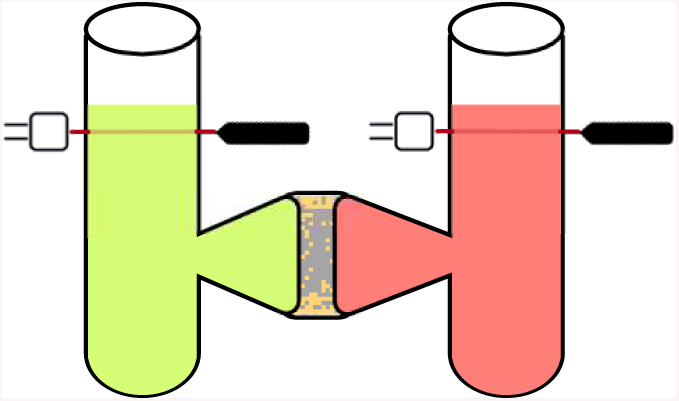
Multicellular Design Idea. Two different chambers are connected through a nanoporose membrane that lets the diffusion of the sensing molecules while blocking the cells. In this way, the two populations can communicate without being mixed.

## ACKNOWLEDGEMENTS

This work was supported by BrisSynBio, a BBSRC/EPSRC Synthetic Biology Research Centre, Grant No. BB/L01386X/1 and by the project COSY-BIO project, which has received funding from the European Union Horizon 2020 programme under Grant Agreement No. 766840.

## Appendix A. SETUP OF THE EXPERIMENT

In addiction to all the references in the above work, our experiments have been conducted with the following setup: Incubator STUART SI 60 D, Syringe Pump and Pinch Valve driven by MG996R High Torque Servos, Syringe size 2.5ml, Air Pump VLIKE silent, 500ml lab bottles for media and waste tanks.

